# Coordination of transporter, cargo, and membrane properties during non-vesicular lipid transport

**DOI:** 10.1101/2023.06.28.546834

**Authors:** Alena Koukalova, Andrea Eisenreichova, Bartosz Różycki, Evzen Boura, Jana Humpolickova

## Abstract

Homeostasis of cellular membranes is maintained by fine-tuning their lipid composition. Yeast lipid transporter Osh6, belonging to the oxysterol-binding protein-related proteins family, was found to participate in the transport of phosphatidylserine (PS). PS synthesized in the endoplasmic reticulum is delivered to the plasma membrane, where it is exchanged for phosphatidylinositol-4 phosphate (PI4P). PI4P provides the driving force for the directed PS transport against its concentration gradient. In this study, we employed an in vitro approach to reconstitute the transport process into the minimalistic system of large unilamellar vesicles to reveal its fundamental biophysical determinants. Our study draws a comprehensive portrait of the interplay between the structure and dynamics of Osh6, the carried cargo lipid, and the physical properties of the involved membranes, with particular attention to their charge and fluidity. Specifically, we address the role of the cargo lipid, which, by occupying the transporter, imposes changes in its dynamics and, consequently, predisposes the cargo to disembark in the correct target membrane.

## Background

Membranes of specific cellular organelles significantly differ in fluidity, thickness and lipid composition [1]. This is an essential feature for their functionality and identity. Most phospholipids are synthesized in the endoplasmic reticulum (ER) and must be transported to their final destination. This process is critical and must be ingeniously orchestrated.

The majority of lipid molecules are transported by vesicles that bud from the source membrane and coalesce with the target membrane [2]. Apart from that, some lipids are also transported via non-vesicular pathways using lipid transfer proteins [3]. The yeast transporter of phosphatidylserine (PS), Osh6 [4-6], belongs to the family of transfer proteins known as oxysterol-binding protein-related proteins (ORPs). Osh6 is a soluble protein localized between the endoplasmic reticulum (ER) and the plasma membrane (PM) [7], where it associates with the membrane tether Ist2 [8]. It transports PS from the ER to the PM by exchanging it for phosphatidylinositol-4-phosphate (PI4P) [9]. By this process, PI4P is released in the ER membrane, where it is dephosphorylated by Sac1 phosphatase [10, 11]. The energy required for the counter-gradient transport of PS is spent on maintaining the PI4P pool in the PM, which consists of the PI transport and its subsequent phosphorylation by phosphatidylinositol-4 kinase.

In our previous work [12], we showed that PS can be transported spontaneously down its gradient without the need for exchange. However, in cells, the transport occurs against the gradient, requiring a deeper mechanistic understanding that includes the different affinities of Osh6 and the individual cargo lipids. PI4P has a higher affinity for the Osh6 binding pocket than PS [4] and can replace the PS molecules, inhibiting the along-gradient transport. To load PS again from the PS-donating membrane, Sac1 is required on that membrane to dephosphorylate the incoming PI4P [12].

Since lipid transport involves not only the interaction between the cargo lipid and the protein, but also the interaction between the entire membrane and the protein, we focus on the role of the membrane in the process. Specifically, we investigate the role of certain membrane features, such as charge and fluidity, on Osh6-mediated lipid transport and relate our findings to the electrostatic properties and dynamics of the transporter. It has been previously shown that Osh6 interacts differently with neutral and charged membranes, and that the interaction is affected by the presence of cargo in the lipid binding pocket [13]. We further elaborate on this by examining the process of cargo extraction and deposition to the target membrane. Our results demonstrate that the kinetics of extraction depend on both the cargo lipid and the characteristics of the entire membrane. Additionally, the cargo lipid influences the interaction of Osh6 with the membrane and determines whether the membrane charge facilitates the release of cargo or not.

## Results

### PS extraction and release

To understand the transport process, we have developed a method that enables us to discriminate between the two components of the transport: i) extraction of the cargo from the donor membrane by Osh6 and ii) release of the cargo to the acceptor membrane. The method employs fluorescence cross-correlation spectroscopy (FCCS) and leverages our understanding of the behavior of the lipid biosensor. We start by preparing large unilamellar vesicles (LUVs) that contain PS and a fluorescent lipid tracer, DiD. Next, we add the PS biosensor, C2 domain of Lactadherin C fused to CFP (C2_Lact_-CFP) [14, 15] to the LUVs. As a result, the LUVs are double-labeled with both CFP and DiD, which leads to a high level of cross-correlation between the CFP and the DiD signal, as both fluorophores move together. Upon addition of Osh6, PS is extracted from the LUVs to the binding pocket of the protein, from where it can be further deposited to a different membrane (another type of LUVs). At this point, the biosensor detaches from the donor membrane, and the cross-correlation amplitude (*G*_cc_(0)) drops down. We can conduct the experiment to either examine the kinetics of the extraction (i) or the release of the cargo to a different membrane (ii).

i) When the level of accessible PS in LUVs is stoichiometrically equivalent to the amount of Osh6, PS extraction from the donor membrane is observed upon Osh6 addition (Fig. 1A,D). The extraction occurs instantaneously and almost irrespective of whether other types of LUVs are present in the system (Fig. 1C). The only exception is for LUVs that contain other Osh6 ligands, such as PI4P, which compromises PS extraction by competing for the binding site. Additionally, PIP_2_ slightly blocks PS extraction, as PIP_2_ [12] has also been shown to be transported by the protein, even though its binding affinity is likely lower compared to the main cargo molecules, such as PS and PI4P.

ii) When the level of accessible PS exceeds the level of Osh6, the addition of Osh6 causes PS extraction, followed by its deposition to the acceptor membrane. However, we can “hide” the extraction in the observation window of our experiment by having so much PS that, even if all of the Osh6 capacity is used, the PS level remains above the dynamic range of the biosensor (Fig. 2A, C). Under these conditions, only a small, fast drop in *G*_cc_(0) is observed upon Osh6 addition when no other LUVs are available (Fig. 2D, E, black squares). However, if acceptor LUVs are present, the drop in *G*_cc_ (0) can be almost solely attributed to ligand release to the acceptor membrane (as the PS extraction is fast) (Fig. 2B, D, E).

**Figure 1.**
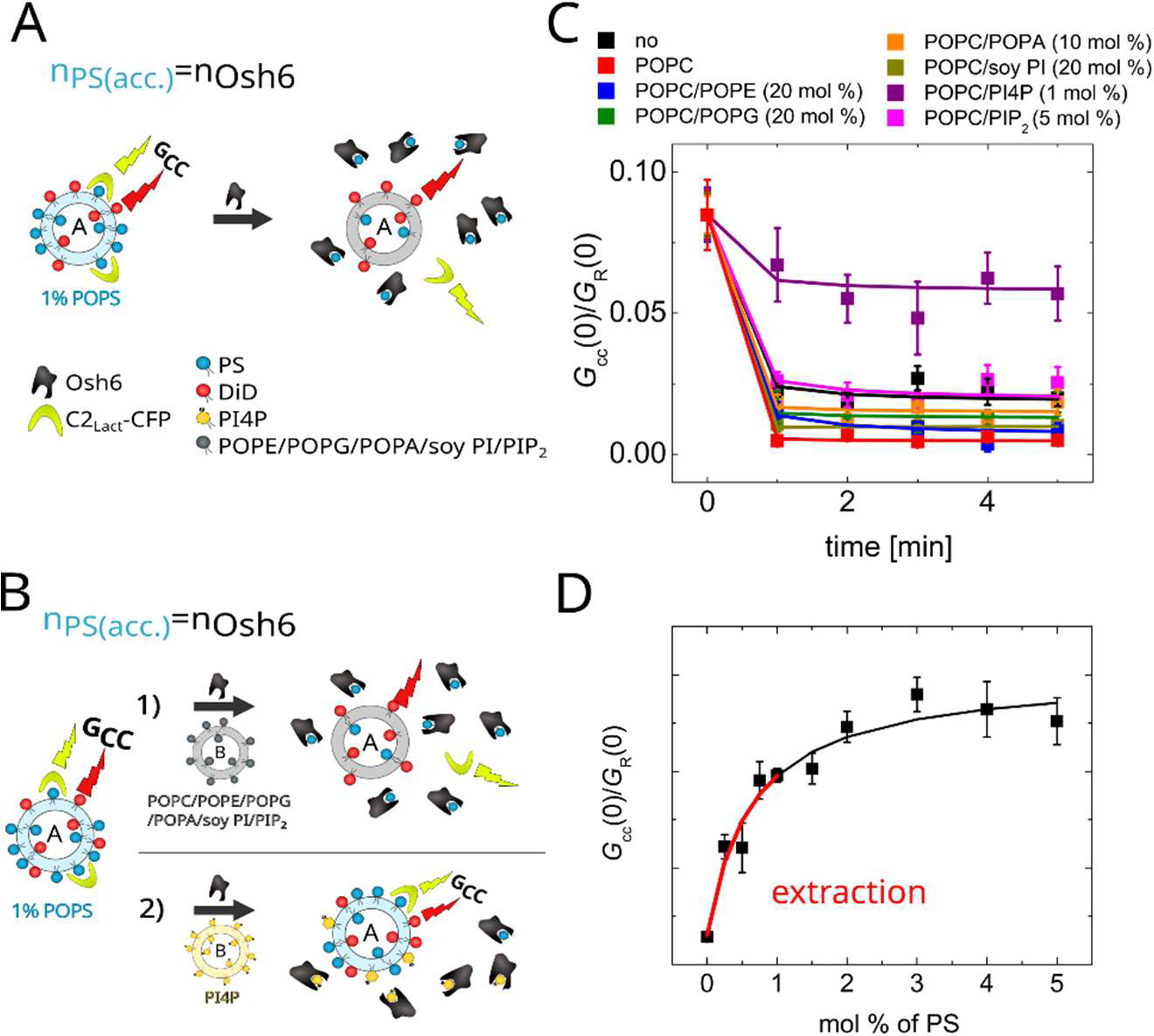
PS extraction assay. A) Scheme of the assay. The amount of accessible PS (*n*_PS_(acc.)) equals the amount of Osh6. The FCCS read-out parameter (*G*_cc_(0)/ *G*_R_(0), simply *G*_cc_) monitoring the mutual motion of DiD and C2-Lact fused to CFP drops upon Osh6 addition. B) Scheme of the assay when the PS donating LUVs are in presence of other, PS-free LUVs either not bearing a competitive ligand (1) – PS extraction occurs, or bearing a competitive ligand (2) – PS extraction is compromised. C) Temporal drop in *G*_cc_ during the PS extraction. The extraction from LUVs containing 1 % PS was carried out in the absence of other, PS-free LUVs (black), or in the presence of other, PS-free LUVs differing in the lipid composition: POPC (red), POPC/POPE (blue), POPC/POPG (green), POPC/POPA (orange), POPC/soy PI (dark yellow), POPC/PI4P (violet), POPC/PIP_2_ (magenta). D) Dependence of the *G*_cc_ read-out of C2_Lact_-CFP biosensor in our experimental system. The red line depicts the region of the calibration curve where extraction can be observed.

**Figure 2.**
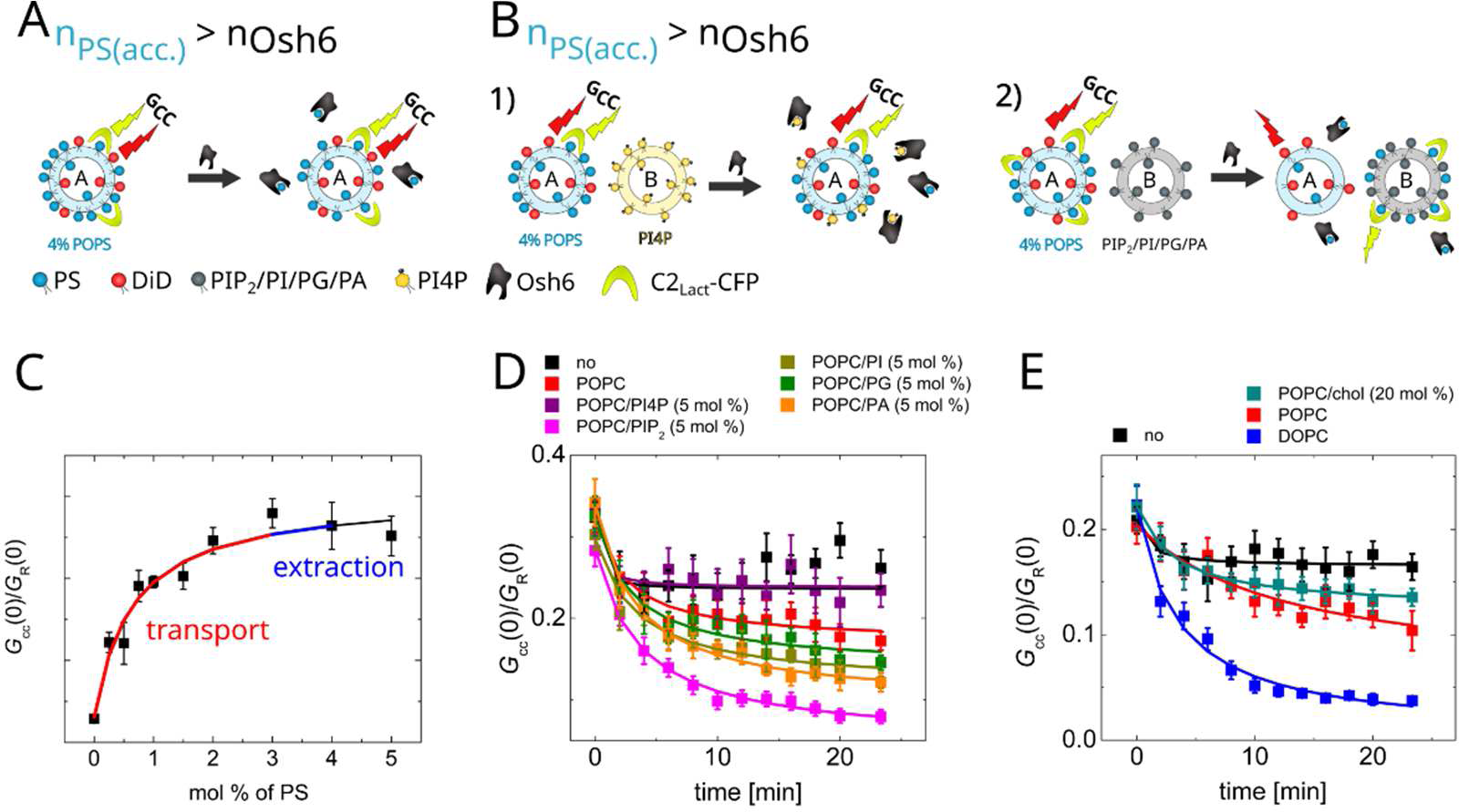
PS transport assay. A) Scheme of the assay. The amount of accessible PS (*n*_PS_(acc.)) is higher than the amount of Osh6. The FCCS read-out, *G*_cc_, monitoring the mutual motion of DiD and C2_Lact_ fused to CFP remains almost unchanged upon the Osh6 addition as the relative change of PS in the donor LUVs is small and the biosensor’s response is saturated. B) Scheme of the assay when the PS donating LUVs are in presence of acceptor LUVs either containing a competitive cargo (1) – Osh6 is blocked by the cargo, or not containing a competitive cargo (2) – transport to the acceptor LUVs occurs. C) Dependence of the *G*_cc_ read-out of C2-Lact biosensor in our experimental system. The blue and red lines show the concentration regions of extraction and transport, respectively. D) Role of charge in the PS release. Temporal drop in *G*_cc_ during the PS transport. No PS accepting LUVs were added (black), PS accepting LUVs were composed of: POPC (red), POPC/PI4P (violet), POPC/PIP_2_ (magenta), POPC/PI (dark yellow), POPC/PG (green), POPC/PA (orange). E) Role of membrane fluidity in the PS release. Temporal drop in *G*_cc_ during the PS transport. No PS accepting LUVs were added (black), PS accepting LUVs were composed of: POPC/cholesterol (cyan), POPC (red), DOPC (blue).

Fig. 2 sheds light on two membrane determinants that significantly affect the release of PS to the accepting membrane: i) charge and ii) membrane fluidity. While the drop in *G*_cc_(0) upon sole extraction of PS from LUVs containing 4 mol % PS is small (Fig. 2D,E black squares), the presence of acceptor LUVs allows for further flow of PS towards the accepting membrane. The extent of the PS release is highly influenced by the charge in the accepting membrane. Lipid head groups of phosphatidylglycerol (PG), phosphatidic acid (PA), and phosphatidylinositol (PI) significantly improve the release compared to the neutral membrane. In contrast, the PI4P head group inhibits PS transport, which is in good agreement with the previous finding that PI4P blocks the binding pocket of Osh6 [12].

We have also paid specific attention to the presence of PIP_2_ in the acceptor membrane, as PIP_2_ was shown to have a great impact on the transport of dehydroergosterol (DHE) by Osh4 [7]. In the case of Osh6-mediated transport of PS, we also observe a significant impact of PIP_2_ on the level of PS transport. However, considering the higher, locally concentrated charge of the PIP_2_ molecule, its effect is comparable to the impact of other charged membranes.

Another important aspect of the PS release to the acceptor membrane is the membrane fluidity. The more fluid the acceptor membrane is, the better it accepts the cargo (Fig. 2E). While cholesterol containing 1-palmitoyl-2-oleoyl-glycero-3-phosphocholine (POPC) almost does not accept the POPS cargo, the membrane composed of highly fluid 1,2-dioleoyl-glycero-3 phosphocholine (DOPC) absorbs the cargo readily.

### PI4P extraction and release

We have shown that PI4P can replace PS in the Osh6 binding pocket and inhibit the along-gradient PS transport. To transport PS against the gradient continuously, PI4P eventually has to be released to the PI4P accepting membrane, i.e., to the ER in cells. We have shown that Sac1-assisted dephosphorylation of PI4P keeps shifting the equilibrium between PI4P bound to Osh6 and PI4P in the accepting membrane [12]. Despite that, before being dephosphorylated, PI4P has to escape the binding pocket while Osh6 gets in contact with the membrane. Here, we focus on the membrane features that facilitate the PI4P release. Similar to our examination of PS, both the PI4P uptake and release are addressed by employing FCCS in a similar setup. The PI4P donating LUVs are labeled with DiD, and PI4P is sensed by SidC-Atto488, so at the beginning, the double-labeled vesicles show high cross-correlation (*G*_cc_) that drops while PI4P is extracted from the donor membrane (Fig. 3A, D).

**Figure 3.**
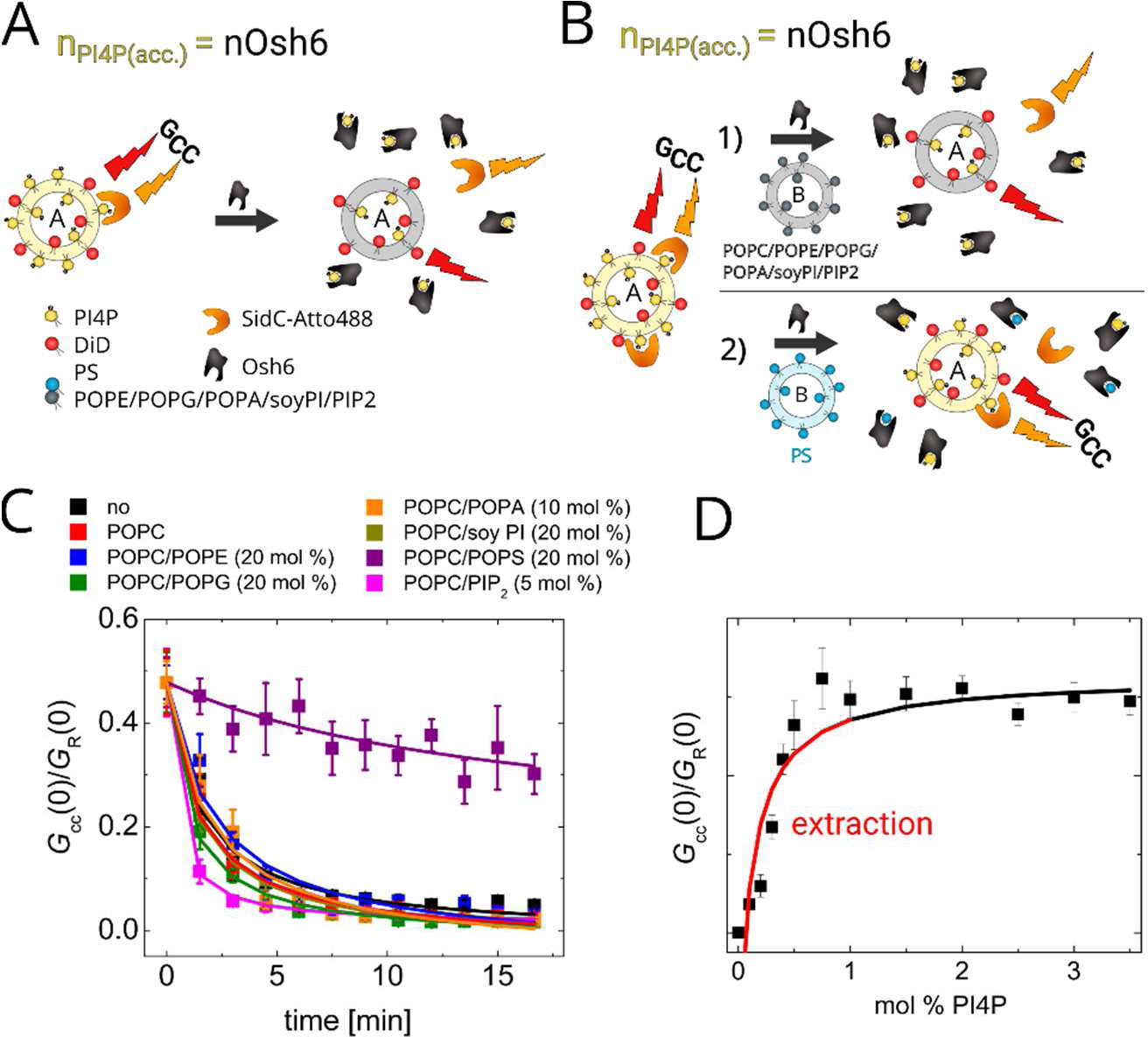
PI4P extraction assay. A) Scheme of the assay. The amount of accessible PI4P (*n*_PI4P_(acc.)) equals the amount of Osh6. The FCCS read-out, *G*_cc_, monitoring the mutual motion of DiD and SidC-Atto488 disappears upon the Osh6 addition. B) Scheme of the assay when other, PI4P-free LUVs (marked as B) are present during the extraction. The LUVs B either do not contain the competing cargo (1) – the extraction is not affected, or contain the competing cargo (2) – the extraction is compromised. C) Kinetics of the PI4P extraction from the PI4P containing LUVs when no other LUVs are present (black), and at presence of other, PI4P-free LUVs of various compositions: POPC (red), POPC/POPE (blue), POPC/POPG (green), POPC/POPA (orange), POPC/soy PI (dark yellow), POPC/POPS (violet), POPC/PIP_2_ (magenta). D) Dependence of the *G*_cc_ read-out of SidC-Atto488 biosensor in our experimental system. Red line depicts the concentration region of PI4P where the extraction takes place.

Fig. 3 addresses the kinetics of PI4P uptake. To achieve this, the level of extractable PI4P is equivalent to the amount of Osh6 used. In addition, the PI4P level is chosen so that the changes in *G*_cc_ dynamically reflect the drop in PI4P in the donor membrane (Fig. 3D). The black curve in Fig. 3C shows that indeed, the entire PI4P can be extracted upon Osh6 addition. It is also worth noticing that PI4P uptake is a much slower process than the uptake of PS. While extracting the PI4P pool requires almost 20 minutes, the kinetics of PS extraction cannot be captured within the temporal resolution of our experiment; all available PS is extracted in less than a minute (compare Fig. 1C and 3C). The presence of other LUVs did not cause significant changes in the PI4P extraction kinetics except for LUVs containing excess PS, which competes with PI4P for the binding pocket (Fig. 3C).

In the following experiments, we focused on the PI4P release to the acceptor membrane. The level of PI4P was set to 3 mol %, *i.e.* above the capacity of Osh6, so that even if all PI4P was extracted, almost no drop in *G*_cc_ was observed as the response of the biosensor under these conditions was saturated (Fig. 4D, black squares).

**Figure 4.**
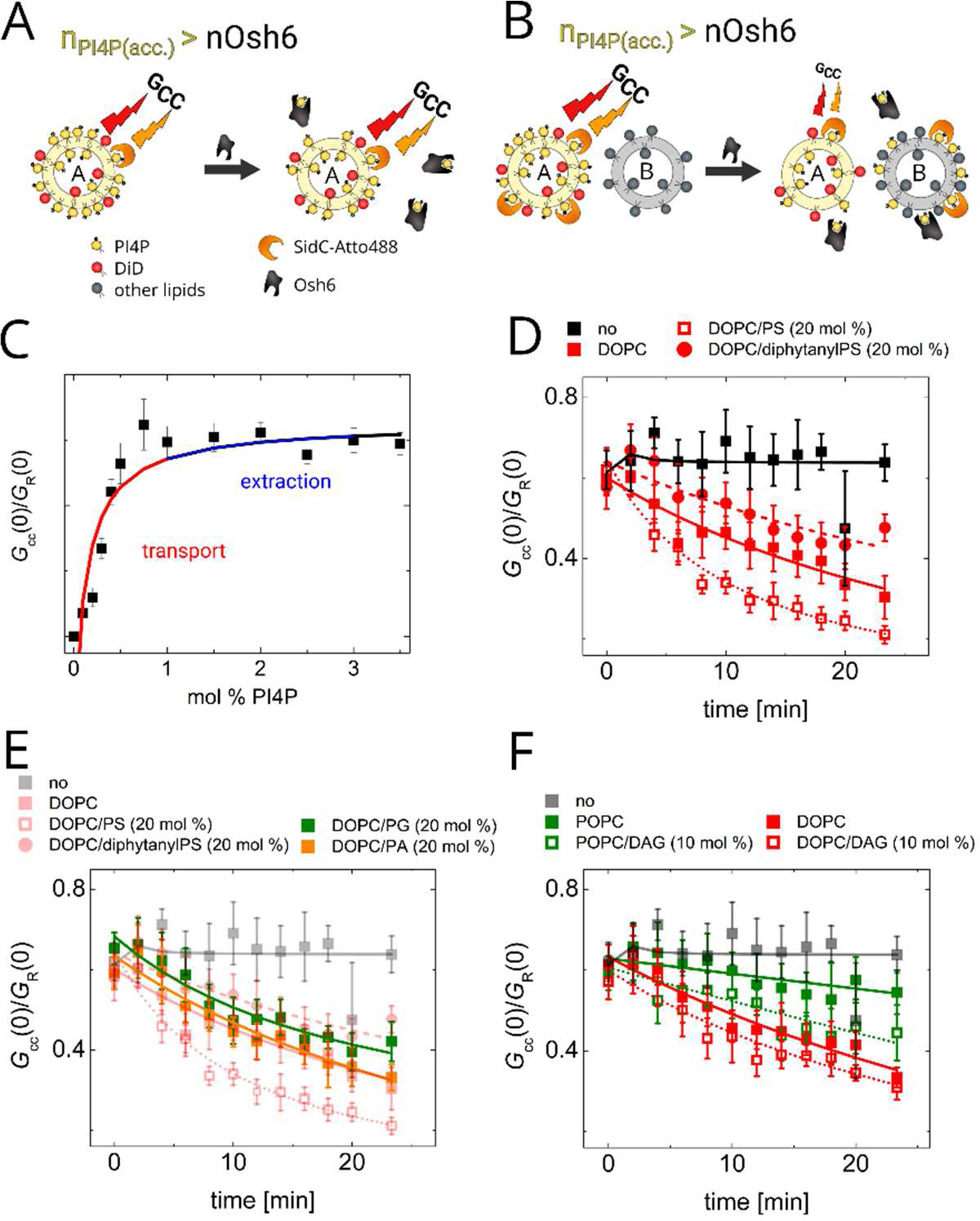
PI4P transport assay. A) Scheme of the assay. The amount of accessible PI4P (*n*_PI4P_(acc.)) exceeds the amount of Osh6. The FCCS read-out, *G*_cc_, does not drop upon the Osh6 addition as the relative change of PI4P level in the PI4P donating LUVs is small and the biosensor’s response is saturated. B) Scheme of the assay when also PI4P accepting LUVs are present in the system. The FCCS read-out, *G*_cc_, drops as the PI4P accepting LUVs allow for the PI4P deposition and continuation of the transport. C) Dependence of the FCCS read-out, *G*_cc_, of SidC-Atto488 biosensor in our experimental system. The blue and red lines depict the PI4P concentration regions where the extraction and the transport can be observed, respectively. D) PI4P transport to LUVs with competing ligand. Kinetics of the PI4P transport to the PI4P accepting LUVs composed of DOPC (red solid squares), DOPC/PS (red hollow squares), DOPC/diphytanoylPS (red solid circles). The experiment with no accepting LUVs is depicted by black solid squares. E) Effect of charge on the PI4P release. The PI4P accepting LUVs were composed of DOPC/PG (green) and DOPC/PA (orange). The data from the D) (partially transparent) are shown for comparison. F) Effect of fluidity on the PI4P release. The PI4P accepting LUVs were composed of POPC (green solid squares), POPC/DAG (green hollow squares), DOPC (red solid squares), and DOPC/DAG (red hollow squares). Experiment without the accepting LUVs is shown for comparison (grey squares).

We then added other LUVs to the system to enable Osh6 to deposit the PI4P cargo to the acceptor membrane. The excess of PS cargo in the acceptor membrane, which can quickly bind to the Osh6 binding pocket, allows for better PI4P release (Fig. 4D, red hollow squares). In contrast, charged membranes that do not contain the cargo (Fig. 4E, orange and green squares) or contain diphytanoylPS (phPS) (Fig. 4D, red solid circles), the lipid with a PS head group and methylated fatty acid chains, which makes it unextractable, reduce the drop of PI4P to the acceptor membrane.

PI4P is obviously better accepted by membranes that are more fluid (DOPC rather than POPC). Also addition of diacylglycerol (DAG) to the PC bilayer promotes PI4P release (Fig. 4F, dotted curves). This is probably due to the fact that DAG induces membrane curvature and by that contributes to better harboring of Osh6 in the acceptor membrane [16].

### Membrane binding of Osh6

To understand the involvement of charge in lipid transfer, a cysteine-less mutant of Osh6 was prepared with a single amino acid residue mutated to cysteine for specific labeling. The transport behavior of the mutant was identical to that of the wild-type protein (data not shown). It was then labeled with Atto488, and its binding to various types of membranes was explored. Osh6-Atto488 was added to LUVs of defined composition that contained DiD, the fluorescence marker. In addition, Osh6-Atto488 was pre-incubated with either PI4P or PS-containing LUVs long enough to saturate the transporter with cargo (Fig. 5A). Using FCCS, the temporal cross-correlation between the green and red signals was monitored.

**Figure 5.**
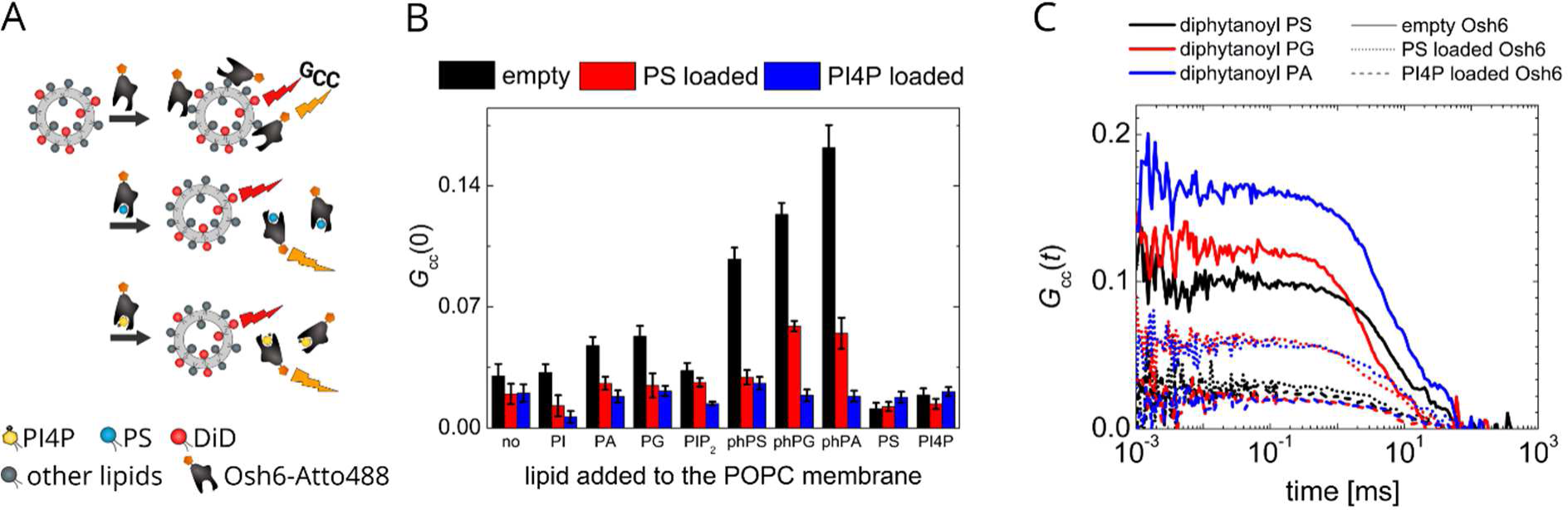
Cargo effect on membrane binding. A) Scheme of the experiment. The DiD labeled LUVs of various compositions were mixed with Atto488-Osh6 that was either empty or pre-incubated with PS- or PI4P-containing LUVs. The binding was observed as the FCCS read-out, *G*_cc_, corresponding to the mutual motion of DiD and Atto488. B) The FCCS read-out, *G*_cc_, monitoring binding Osh6-Atto488 to membranes of various lipid composition of depending (POPC membrane contained 5 mol % of PI4P or PIP_2_ and 20 mol % of other charged lipids). Black column stays for the empty protein, red for the PS-loaded, and blue for the PI4P-loaded one. C) FCCS temporal crosscorrelation curves of Osh6-Atto488 and DiD-labeled LUVs of different composition: POPC + 20 mol % diphytanoylPS (black), POPC + 20 mol % diphytanoylPG (red), POPC + 20 mol % diphytanoylPA (blue). The added Osh6 was either empty (solid lines), or pre-incubated with PS-containing unlabeled LUVs (dotted lines), or with PI4P-containing unlabeled LUVs (dashed lines).

Fig. 5B shows the amplitudes of the cross-correlation curves between the Osh6-Atto488 and the DiD signals, which relate to the interaction between the protein and the membrane. The higher the amplitude is, the more labeled Osh6 molecules are attached to the membrane. First, the interaction of the protein with various LUVs was monitored (Fig. 5B, black columns). Keeping in mind that PS and PI4P are the main cargo lipids, and that the membrane containing these lipids will also provide Osh6 with the cargo, we can conclude, in agreement with the literature [13], that Osh6 binds weakly to the cargo-donating membrane. Slightly more pronounced adhesion to the neutral membrane is observed. Osh6 adheres better to membranes that contain charged lipids such as PG and PA and best to membranes containing lipids with negatively charged headgroups and with phytanoyl chains, such as diphytanoylPG (phPG), diphytanoylPA (phPA), or diphytanoylPS (phPS). Osh6 recognizes phPS as a ligand but, due to the methylated lipid chains, cannot accommodate it and pull it out from the membrane. phPG and phPA do not contain a ligand headgroup, yet both serve as a platform for Osh6 binding. The difference in the Osh6 binding between the phytanoyl tails in phPG and phPA and the palmitoyl and oleoyl chains in POPG and POPA, respectively, can perhaps be attributed to better exposure of the charge due to lipid packing defects [17]. The binding is not much affected by the charge of PI, which may be shielded by the inositol ring. Also, PIP_2_ is a highly negatively charged lipid that does not significantly increase the membrane adhesion of Osh6, but it has to be kept in mind that PIP_2_ was also identified as a weakly binding cargo of Osh6 [12].

Second, to learn about the interaction of PS- or PI4P-loaded Osh6, we pre-incubated Osh6-Atto488 with label-free LUVs containing either PS or PI4P. We then added the mixture to the solution of labeled LUVs (Fig. 5B, red and blue columns). Adhesion to all types of labeled cargo-free LUVs dropped, which was more pronounced for the PI4P-loaded Osh6. Fig. 5C shows the temporal cross-correlation curves for the LUVs containing lipids with phytanoyl tails: phPS (black lines), phPG (red lines), and phPA (blue lines). The solid, dotted, and dashed lines show the interaction of empty, PS-, and PI4P-loaded Osh6, respectively.

Jointly, these results clearly suggest that the cargo changes the protein-membrane interaction, specifically, that the PS release from the binding pocket is charge-dependent compared to the PI4P release.

### Role of the cargo in membrane distinguishing

As we have seen already, the cargo loaded protein interacts differently with various types of membranes depending on the cargo it is loaded with (Fig. 5). Also, the knowledge on the extraction kinetics raises questions dealing with the situation in cells. For example, it is not clear how PI4P gets extracted from the PS rich plasma membrane if the PS extraction is a much faster process than the extraction of PI4P. Motivated by this discrepancy, we examined the extraction of PI4P from highly charged membranes using the assay shown in Fig. 4, paying particular attention to the difference between the empty protein and the protein doped with PS.

Fig. 6B shows that from the PG-rich membrane (20 mol % POPG), PI4P can be extracted much faster than from the neutral one. A different observation was made for 20 mol % POPS in the membrane, as PS is also the cargo molecule. In this case, the extraction of PI4P is impaired. The scenario, however, changes dramatically if the protein already contains the PS cargo (the protein was incubated with the PS LUVs prior to the addition). In that case, PI4P gets extracted readily. Additionally, PS-loaded Osh6 does not extract PI4P from the neutral membrane. This suggests that to be extracted, PI4P requires (i) a charged membrane and (ii) PS-loaded Osh6.

**Figure 6.**
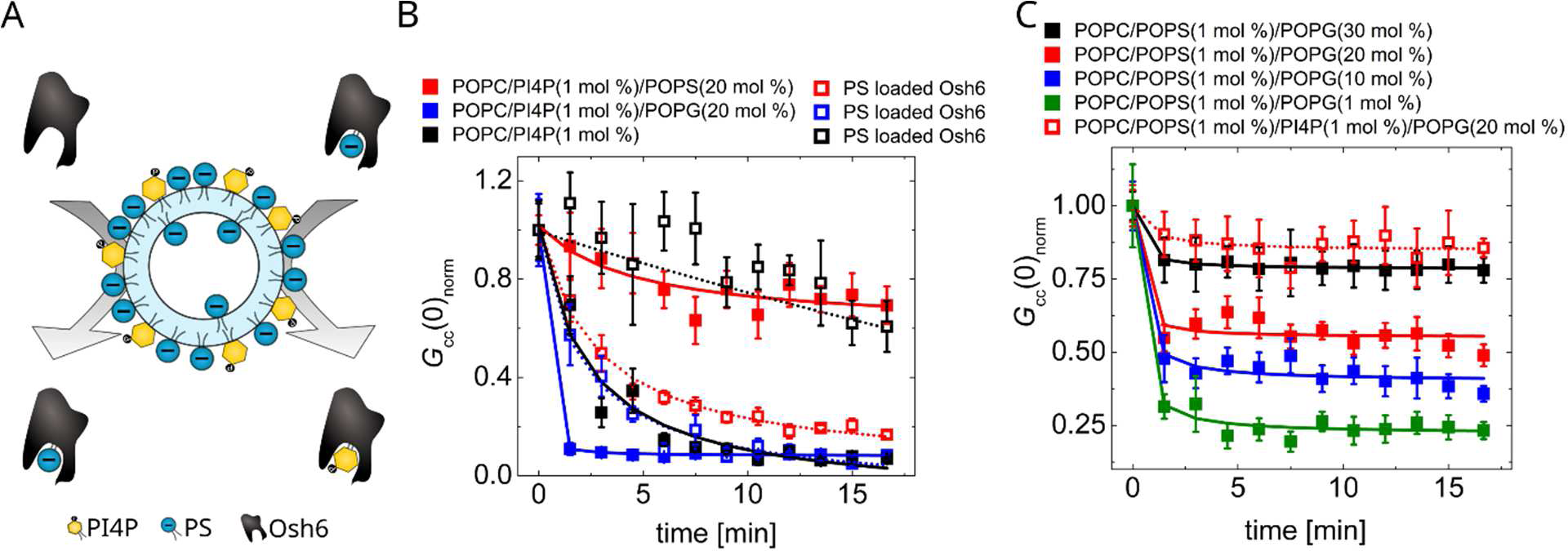
Synchronization of charge and cargo occupancy required for the PI4P extraction from the PM-like membrane. A) Schematic description of the PI4P extraction experiment: PI4P is only extracted from the PM-like membrane (rich in PS), if Osh6 that approaches the membrane is PS-loaded. B) PI4P extraction assay. Temporal dependence of the normalized FCCS read-out, *G*_cc_(0)_norm_, monitoring the PI4P drop upon Osh6 addition. PI4P is extracted from various kinds of LUVs: POPC/PI4P (1 mol %)/POPS (20 mol %) (red squares), POPC/PI4P (1 mol %)/POPG (20 mol %) (blue squares), and POPC/PI4P (1 mol %) (black squares). The solid and the hollow squares stand for empty and PS-loaded Osh6 addition, respectively. C) Charge-dependent PS extraction kinetics. Temporal dependence of the normalized FCCS read-out, *G*_cc_(0)_norm_, monitoring the PS drop upon Osh6 addition. The LUVs for the PS extraction (1 mol % of POPS) contained gradually increasing amount of negatively charged POPG: 1 mol % (green squares), 10 mol % (blue squares), 20 mol % (red squares), 30 mol % (black squares). Red hollow squares depict the PS extraction kinetics from LUVs that except from 20 mol % of POPG contained additional 1 mol % of PI4P.

Next, we examined whether the PS-loaded Osh6 requires PS in the PI4P-containing membrane to extract it, or whether non-specific charge would be sufficient. We have prepared a PI4P bilayer containing only 1.5 % of PS (instead of 20 %) and repeated the experiment first with the empty Osh6. The presence of PS competes with the PI4P extraction as shown on the Fig. S1. With PS-loaded Osh6, the PI4P extraction is even more impaired. To discern the effect of high charge from the specific effect of high PS, we have completed the 20 % of charged lipid not with PS, but with 18.5 % of PG. In this case, PI4P was again well recognized and extracted by PS-loaded Osh6 (Fig. S1).

Altogether, this report highlights an important mechanistic determinant of transport, namely, that Osh6 can only extract PI4P from the plasma membrane (a membrane that is highly charged) when it contains the PS cargo.

### Role of the charge in the PS extraction

We have already shown that the rate of PI4P extraction is charge-dependent. In highly charged membranes, such as the plasma membrane, PI4P is extracted faster than from neutral membranes. To complete the picture, we examined the kinetics of PS extraction from charged membranes. We extracted PS from charged membranes with increasing amounts of POPG (5 to 30 %, Fig. 6D). The higher the amount of PG (negative charge) in the membranes, the lower the fraction of PS that can be extracted, with a significantly reduced rate. When the membrane composition is enriched with PI4P in an equimolar ratio to PS, the extraction of PS is almost entirely compromised.

### The cargo dependent protein dynamics

To get insights into molecular details of Osh6 motions, we performed all-atom molecular dynamics (MD) simulations of the POPS-loaded and PI4P-loaded Osh6 in solution. For each of the two systems, we conducted two simulations of 500 ns each, resulting in a total of 2 µs of MD data for analysis. To quantify segmental motions of Osh6 with the two cargo lipids, we computed the root-mean-square fluctuations (RMSF) along the Osh6 primary structure (Fig. 7A). We found that the N-terminal segment comprising amino acid residues with numbers from 35 to 48 was displaced and quite mobile in the POPS-loaded Osh6, while it was clearly less mobile in the PI4P-loaded Osh6. Also the tip of helix 11 comprising amino acid residues with numbers from about 360 to 370 was found to be more mobile in the POPS-loaded Osh6 than in the PI4P-loaded Osh6. On the other hand, the C-terminal segment comprising amino acid residues with numbers from 420 to 435 was found to be more mobile in the PI4P-loaded Osh6 than in the POPS-loaded Osh6. We also noticed that the loops that were most flexible were those comprising amino acid residues with numbers from 206 to 211 and from 256 to 262. They were more-or-less equally mobile in each of the Osh6 load states in our MD simulations.

**Figure 7.**
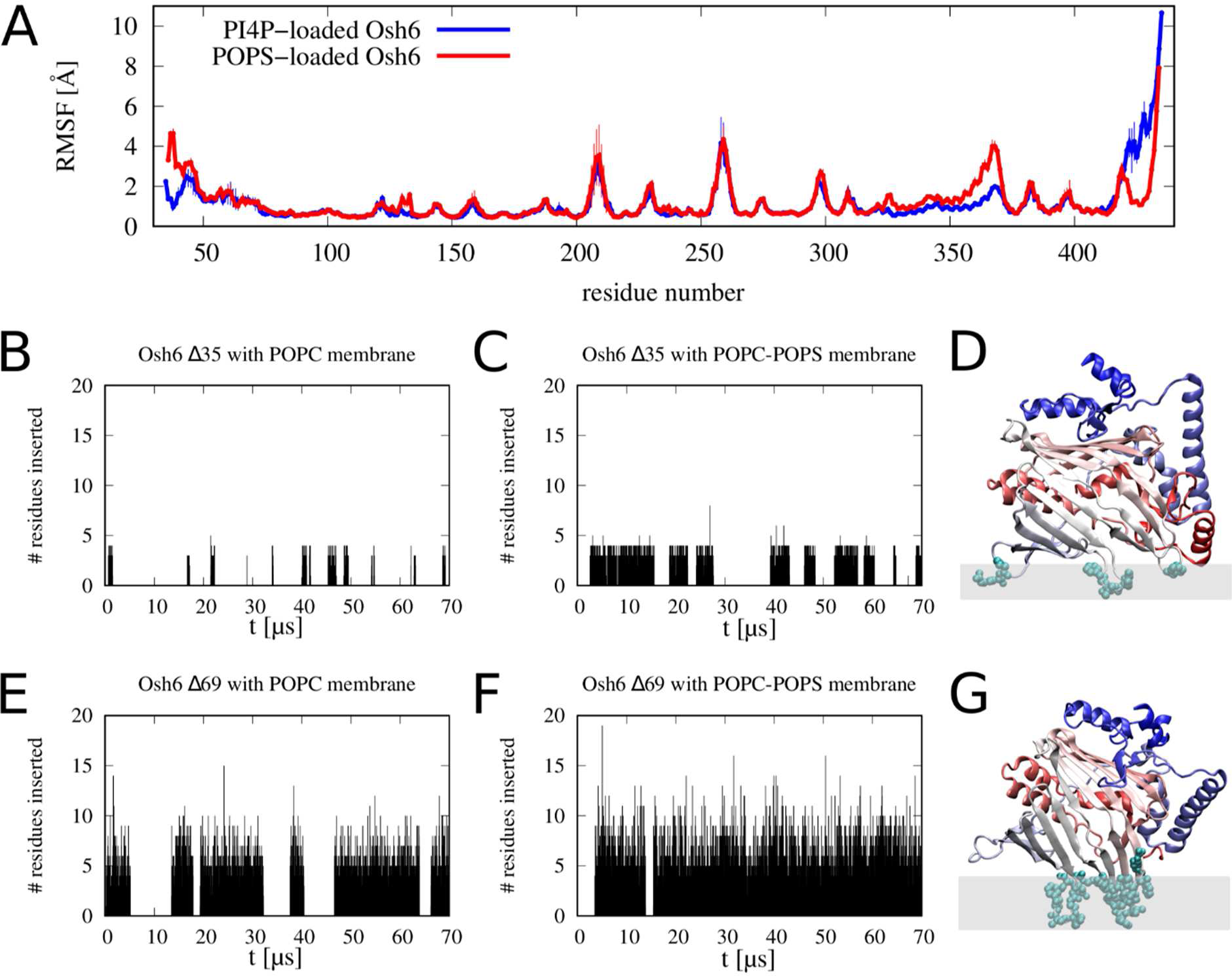
MD simulation results. (A) RMSFs obtained from the all-atom MD simulations of the POPS-loaded (red) and PI4P-loaded (blue) Osh6 in solution. (B,C) Number of amino acid residues inserted into the POPC (B) and POPC-POPS (C) bilayer as a function of time. Results obtained from the coarse-grained MD simulations of Δ35. (D) Cartoon illustrating how Δ35 associates with lipid membranes. Δ35 is colored red-to-blue (from the N-terminus to the C-terminus). The amino acid residues that get inserted into the lipid bilayer are shown in the van der Waals representation. (E,F) Analogous to panels B and C but for Δ69. (G) Analogous to panel D but for Δ69.

As already emphasized by Lipp *et al* [13], the N-terminal lid contains the D/E-rich motif that compensates for the basic surface of Osh6 that is below the lid. Lipp therefore considers the Δ69 mutant of Osh6 as an “open” form of Osh6. The MD simulations showing larger flexibility of the N-terminal segment in the case of the PS-loaded Osh6 suggest that the PS-loaded transporter is more likely to open and to associate with the negatively charged surface compared to the PI4P-loaded one.

In order to study how Osh6 interacts with lipid bilayers on the timescale of the order of 100 µs, we performed coarse-grained MD simulations using the Martini 3 model [18]. We simulated four systems: 1) Δ35, the “closed” form in contact with a POPC bilayer, 2) Δ35 in contact with a charged bilayer comprising POPC and POPS in 4:1 molar ratio, 3) Δ69, the “open” form in contact with the POPC bilayer, and 4) Δ69 in contact with the POPC-POPS bilayer. For each of the four systems, we monitored the number of Osh6 amino acid residues inserted into the lipid bilayer as a function of time (Fig. 7 B,C,E,F). Here, we defined an amino acid residue to be inserted into a bilayer at a given time instance if any of its atoms were in a distance smaller than 1.5 nm from the bilayer midplane. The data in Fig. 7 B-G demonstrate that Δ69 associates with lipid bilayers more strongly than Δ35, and the electrostatic charge on the bilayer enhances the association of both Δ69 and Δ35. We also analyzed which amino acid residues were inserted into the lipid bilayer in the course of the simulations. In the case of Δ35, these were PHE 229, TYR 258, VAL 259, PHE 260, PRO 297, and ARG 298 (Fig. 7D), localized within three flexible loops. In the case of Δ69, the membrane insertion site was found to be significantly larger and formed by three segments: from ARG 184 to SER 189, from ARG 225 to ARG 233, and from LYS 256 to TYR 263 (Fig. 7G). Thus the orientations of Δ35 and Δ69 at the lipid bilayer were different.

## Discussion

In this study, we have conducted a comprehensive biophysical analysis of the lipid transfer protein Osh6 and its role in lipid transportation. While we acknowledge that other biological factors, such as regulatory proteins, energy-loaded pools of PI4P, and membrane tethers, also play crucial roles in this process, the fundamental biophysical interplay between the transporter’s structure, its cargo, and the characteristics of the involved membranes serve as the underlying mechanistic platform of this process.

Most of the studies focus on the Osh6 transporter and its cargo molecules of various head groups and fatty acyl chains [12, 19, 20] without addressing the milieu accommodating the cargo – the membranes. In the case of Osh4, an ergosterol transporter, the saturated fatty acyl chains in the sterol accepting membrane help stabilize the ergosterol gradient that needs to be established between the membrane of trans-Golgi and ER [9]. Additionally, the interaction between the distal membrane binding site of Osh4 and PIP2 has been identified to play a part in the sterol transport [7]. Thus, the molecular view of the transport dynamics does not only involve the protein/ligand interactions but also the protein/membrane, as well as the cargo lipid–ligand/membrane interactions. Moreover, in addition to the thermodynamic affinities between the key players, it turns out that the kinetics of the individual transport steps also come into play.

To ensure the biological relevance of our investigation, we pose questions that are significant for PS transport in cell-like membranes. One intriguing question is related to the fact that PM contains more than 20 mol % of PS, predominantly in its inner leaflet, compared to the low level of PS in ER [1]. Therefore, PS transport needs to occur against this large concentration gradient. Even though PI4P is a stronger binder of Osh6 [4], statistically, the exchange of PS delivered by Osh6 from ER at the PM surface would favor the far more abundant molecule – PS. Additionally, our findings demonstrate that the PS extraction from the neutral membrane is significantly faster than that of PI4P. The delivery of PS to the PS-rich membrane seems to be feasible only if: i) the membrane contains PI4P for the exchange, and ii) the membrane is highly charged, which is true for PM. It should be noted that while PS can be spontaneously and independently delivered along the gradient, transport against the gradient requires high charge and PI4P for the exchange. The highly charged membrane promotes the release of PS, accelerates the uptake of PI4P, and inhibits the non-effective re-extraction of PS. From the perspective of the cargo, PS-loaded Osh6 is poised to be unloaded at the PM surface.

The other question to be answered concerns PI4P offloading. Our experiments show that, contrary to PS, PI4P disembarkation is not enhanced by membrane charge. Generally, in line with the higher affinity of Osh6 for PI4P compared to PS, PI4P-loaded Osh6 is significantly more retentive. The release is enhanced in soft, low-charged membranes that are endowed by phosphatase to shift the equilibrium [10, 12], such as Sac1 in the ER membrane.

Examination of the molecular dynamics of PS- and PI4P-loaded Osh6 in solution by means of all-atom MD simulations revealed a larger mobility of the N-terminal lid in the case of PS-loaded Osh6. As already suggested by others [13], the lid containing the D/E-rich motives shields a basic patch of the protein surface. This suggests that the enhancement of the PS-release by the negatively charged membranes reflects the involvement of the basic amino acid residues of the semi-open Osh6 structure. In the case of the PI4P-loaded Osh6, the lid is firmly closed, and thus PI4P is released in a charge-independent manner.

To complement the all-atom simulations of Osh6 in solution, Martini coarse-grained simulations of the protein were performed to examine its interactions with charged and neutral membranes. The lidless Δ69 mutant represented the “open” form of the protein, and the Δ35 mutant was considered the “closed” one. The “open” form showed a significantly stronger association with the membrane, especially when negatively charged. This finding additionally confirms that our ∼50 µs Martini simulations provide predictions consistent with those obtained from the ∼500 ns all-atom simulations of Lipp *et al* [13].

In accordance with the work of Lipp *et al.*, our findings confirm that the membrane interaction of Osh6 is reduced when the membrane contains the cargo, and the reduction is closely related to the status of the N-terminal lid. In our work, we furthermore claim that the type of cargo governs the behavior of the protein’s action at the specific moment of the PS transport cycle, *i.e.*, PS- and PI4P-loaded Osh6 is determined to offload the cargo in PM and ER, respectively.

## Conclusions

Our study provides a detailed insight into the biophysical orchestration of lipid transport by the lipid transfer protein Osh6. We demonstrate, for the first time, that the cargo itself induces changes in the protein dynamics, specifically its propensity to open its lid. This influences the transporter’s response towards the appropriate target membrane. While carrying PS loosens the lid and exposes the transporter’s basic surface to release the cargo into highly charged membranes (such as the PM), carrying PI4P tightens the lid, necessitating a low-charged, fluid membrane (such as the ER) to accommodate the PI4P unloading.

## Material and Methods

### Protein expression and purification

The genes encoding Osh6, SidC, and C2_Lact_-CFP were cloned into a modified pHIS2 vector containing an N-terminal His6x-tag followed by a tobacco etch virus (TEV) cleavage site. Mutations to produce cysteine-less protein (C62S/C162S/C389S) and a single cysteine mutant (I241C) were generated using site-directed mutagenesis. The proteins were expressed in E. coli BL21 Star cells using our standard protocols [21, 22]. The cells were harvested by centrifugation, resuspended in a lysis buffer (50 mM Tris, pH = 8, 300 mM NaCl, 20 mM imidazole, 3 mM β-mercaptoethanol, and 10% glycerol), and lysed by sonication. The lysates were cleared by centrifugation and incubated with nickel-charged affinity resin (Machery-Nagel). The proteins were eluted with an elution buffer (lysis buffer supplemented with 300 mM imidazole), and the His6x-tag was cleaved off by TEV protease.

Subsequently, the proteins were purified by size exclusion chromatography on a HiLoad 16/600 Superdex75 pg column (Cytiva) in 20 mM Tris, pH 7.4, 300 mM NaCl, 3 mM β-mercaptoethanol, and 10% glycerol. Osh6 was further purified by ion-exchange chromatography on a HiTrap SPHP column (Cytiva).

For Atto488 labeling, the single cysteine mutant of Osh6 and SidC were transferred to PBS and mixed with a 3-fold molar excess of Atto488 maleimide. After overnight incubation at 4 °C, the unbound dye was removed by size exclusion chromatography on Sup200 in 20 mM Tris, pH 7.4, 300 mM NaCl, 3 mM β-mercaptoethanol, and 10% glycerol.

### Lipids and other chemicals

All lipids were purchased from Avanti Polar Lipids (Alabaster, AL) and were used as obtained. Atto488-labeled DOPE was obtained from ATTO-TEC (Siegen, Germany), and the lipid tracer DiD and other basic chemicals were purchased from Sigma-Aldrich (St. Louis, MO).

### LUV formation

LUVs were prepared by extrusion described elsewhere [23]. Briefly, lipids in organic solvents were mixed in the desired ratio so that the final lipid concentration in LUVs was 1 mM. Organic solvents were evaporated in a stream of nitrogen and kept in the vacuum chamber for at least one hour. Later, the lipid films were resuspended in the LUV buffer (40 mM imidazole (pH = 7.4), 150 mM NaCl, 3 mM beta-mercaptoethanol, 1 mM EDTA), and 50 nm LUVs in diameter were prepared in the extruder through the membrane of an appropriate pore size.

### Kinetic assays

All the kinetics was acquired as a short 200-second measurement prior the transporter addition and a longer (5 to 20 minute) measurement upon its addition. The Osh6 concentration was 250 nM. The concentration of the biosensors C2_Lact_-CFP and SidC-Atto488 in the total volume of 200 μL was 50 and 100 nM, respectively.

### PS extraction assays

10 µl of donor LUVs containing POPC (94 mol %), diphytanoy-PG (5 mol %) and POPS (1 mol %) labelled with DiD were mixed with C2_Lact_-CFP, the LUV buffer and 0 µl, or 40 µl of unlabelled LUVs composed of POPC and various contents of other lipid species, as indicated in related figures (Fig. 1C, 6C).

### PS transport assays

10 µl of donor LUVs containing POPC (91 mol %), diphytanoyl-PG (5 mol %), and POPS (4 mol %) labeled with DiD was mixed with C2_Lact_-CFP, the LUV buffer and 0 µl or 40 µl of unlabelled LUVs of various lipid compositions as indicated in Fig. 2D or Fig. 2E.

### PI4P extraction assays

10 µl of donor LUVs containing POPC (99 mol %) and PI4P (1 mol %) labelled with DiD were mixed with SidC-Atto488, the LUV buffer, and 0 µl or 40 µl of unlabelled LUVs composed of POPC and various amounts of other lipids, as indicated in Fig. 3C.

In the experiments that required loading of Osh6 with PS (Fig 6B), Osh6 was pre-incubated with 20 µl of unlabelled LUVs composed of POPC/POPS (80/20 mol %) for 10 min. Next, 20 µl of DiD labelled LUVs composed of POPC/PI4P (99/1 mol %) or POPC/PI4P/POPS (79/1/20 mol %) was mixed with the LUV buffer and SidC-Atto488. Subsequently, a short 200-second measurement was started. The pre-incubated sample of Osh6 and POPC/POPS LUVs were then added, and immediately after that, another 15-minute FCCS measurement was started. The concentration of Osh6 in the final 200 µl sample was 250 nM, and the concentration of SidC-Atto488 was 100 nM.

### PI4P transport assays

10 µl of donor LUVs containing POPC (97 mol %), and PI4P (3 mol %) labelled with DiD were mixed with SidC-Atto488, the LUV buffer, and 0 µl or 40 µl of unlabelled LUVs composed of lipid mixtures, as indicated in Fig. 4D or 4F.

### Osh6 membrane binding

Osh6-Atto488 was incubated with unlabelled LUVs composed of DOPC/POPS (80/20) or DOPC/PI4P (95/5) for 10 min. Subsequently, these samples were incubated with DiD-labelled LUVs of different compositions specified in Fig. 5B for 10 minutes, and a 200-second FCCS measurement was carried out. In the case of empty Osh6, Osh6-Atto488 was added directly to DiD-labelled LUVs. The final concentration of Osh6-Atto488 in all samples was 10 nM.

The labeling of LUVs with DiD was done in the DiD/lipid = 1/10000 ratio. All the FCCS experiments were acquired at least in three independent experiments to ensure for reproducibility.

### Microscopy

The FCCS experiments were carried out on the Leica SP8 confocal microscope (Leica, Mannheim, Germany) equipped with a high numerical aperture water objective (63x, N.A. = 1.2), a battery of synchronizable pulsed lasers, and sensitive hybrid HyD detectors. In our experiments, we used the 640 nm line of the white light laser (Coherent, Inc., Santa Clara, CA) for DiD excitation and the 440 nm and 470 nm diode laser heads (Picoquant, Berlin, Germany) for CFP and Atto488 excitation, respectively. The pair of lasers (440/640 and 470/640) was pulsing alternatively at the pulsed interleaved excitation (PIE) mode at an overall repetition frequency of 40 and 20 MHz, respectively. The PIE mode was used to apply temporal filtering of photon arrival times in addition to the spectral information to omit bleed-through. The data were correlated and evaluated by home-written scripts in Matlab (Mathworks, Natick, MA).

### All-atom MD simulations

The atomic coordinates of the POPS-loaded and PI4P-loaded Osh6 were taken from the crystal structures deposited in the Protein Data Bank (PDB) with the entry codes of 4B2Z and 4PH7, respectively [4, 5]. Systems for MD simulations were prepared using the input generator on the CHARMM-GUI website [24, 25]. Each of the protein structures was solvated in a cubic box with the side length of 9 nm. Sodium and chloride ions were added to neutralize the systems and to reach a physiological ion concentration of 150 mM. The initial systems for MD simulations were energy-minimized using a conjugate gradient method and then equilibrated in a standard procedure using input files provided by the CHARMM-GUI input generator.

The MD simulations were performed using NAMD 2.14 with CHARMM36 force field and the TIP3P model for water molecules [26-29]. Temperature was kept at 303 K through a Langevin thermostat with a damping coefficient of 1/ps. Pressure was maintained at 1 atm using the Langevin piston Nose-Hoover method with a damping timescale of 25 fs and an oscillation period of 50 fs. Short-range nonbonded interactions were cutoff smoothly between 1 and 1.2 nm. Long-range electrostatic interactions were computed using the particle mesh Ewald method with a grid spacing of 0.1 nm. Simulations were performed with an integration time step of 2 fs.

For each of the simulation systems (i.e. the POPS- and PI4P-loaded Osh6) we performed two production runs of 500 ns each, amounting to 2 µs of MD data for analysis. The simulation trajectories were visualized and analyzed using VMD [30].

### Coarse-grained MD simulations

Two variants of Osh6 in contact with lipid membranes were simulated using the Martini 3 model: Δ35 (comprising amino acid residues with numbers from 36 to 434) and Δ69 (comprising amino acid residues with numbers from 70 to 434). The atomic coordinates of Δ35 and Δ69 were taken from the crystal structures deposited in the PDB with the entry codes of 4PH7 and 4B2Z, respectively [4, 5].

Systems for coarse-grained MD simulations were set up in the following way using the Martini maker on the CHARMM-GUI input generator website [24, 31]: Two bilayer segments with equal lateral dimensions of 12 nm by 12 nm were formed independently. In one case, the bilayer was composed of 352 POPC lipids and 88 POPS lipids (i.e. with 4:1 molar ratio). In the other case, the bilayer was composed of 422 POPC lipids. Then each of the two Osh6 structures was placed in a random orientation about 4 nm above each of the lipid bilayers, which produced four simulation systems, i.e., Δ35 with the POPC-POPS bilayer, Δ35 with the POPC bilayer, Δ69 with the POPC-POPS bilayer, and Δ69 with the POPC bilayer. Each of these systems was placed in a cuboid box and solvated. Sodium and chloride ions were added to neutralize the systems and to reach a physiological ion concentration of 150 mM. The simulation systems were coarse-grained within the framework of the Martini 3 model with an elastic network (ELNEDIN) applied to Osh6 beads [18, 32]. The initial systems for MD simulations were energy-minimized using a conjugate gradient method, and then equilibrated in a standard procedure using input files generated by the Martini maker in the CHARMM-GUI input generator.

The coarse-grained MD simulations were performed using Gromacs 2020.2 and the Martini 3.0 force field [18, 33, 34]. The temperature and pressure were kept constant at T = 303 K and p = 1 bar, respectively, using the velocity-rescaling thermostat and the Parrinello-Rahman barostat [35, 36]. Nonbonded interactions were treated with the Verlet cutoff scheme. The cutoff for van der Waals interactions was set to 1.1 nm. Coulomb interactions were treated using the reaction-field method with a cutoff of 1.1 nm and dielectric constant of 15. The integration time step was set to 20 fs. For each of the four systems we performed a MD simulation run of 70 µs. Frames were saved every 1 ns. The simulation trajectories were post-processed with MDVWhole to treat the periodic boundary conditions, and visualized using VMD [30].

## List of abbreviations

ER: endoplasmic reticulum
PM: plasma membrane
PS: phosphatidylserine
PI4P: phosphatidylinositol-4 phosphate
PIP_2_: phosphatidylinositol-4,5 bisphosphate
PI: phosphatidylinositol
POPC: palmitoyl-oleoyl-phosphatidylcholine
POPS: palmitoyl-oleoyl-phosphatidylserine
DOPC: dioleoyl phosphatidylcholine
POPG: palmitoyl-oleoyl-phosphatidylglycerol
POPA: palmitoyl-oleoyl-phosphatidic acid
DAG: diacyl-glycerol
phPS: diphytanoyl-phosphatidylserine
phPG: diphytanoyl-phosphatidylglycerol
phPA: diphytanoyl-phosphatidic acid
ORP: oxysterol-binding protein related protein
LUV: large unilamellar vesicle
FCCS: fluorescence crosscorrelation spectroscopy
RMSF: root mean square fluctuation
C2_Lact_: C2 domain of lactadherin

## Declarations

## Ethics approval and consent to participate

Not applicable.

### Consent for publication

Not applicable.

### Availability of data and materials

All data generated or analyzed during this study are included in this published article and its supplementary information files.

### Author contributions

AK and JH designed the experiments. AK conducted the FCCS measurements, while JH evaluated the data. AE prepared, purified, and labeled the proteins. BR carried out the simulations. EB supervised the project. JH wrote the manuscript, and all contributors participated in the data interpretation and manuscript preparation.

### Funding

The research was supported jointly by the Czech Science Foundation, grant number 21-27735K (to JH and EB), and by the National Science Centre, Poland, grant number 2020/02/Y/NZ1/00020 (to BR), within the international CEUS-UNISONO program. The molecular dynamics simulations were performed using the supercomputer resources at the Centre of Informatics -- Tricity Academic Supercomputer and networK (CI TASK) in Gdansk, Poland. We also acknowledge the Academy of Sciences of the Czech Republic (RVO: 61388963).

### Author details

^1^Institute of Organic Chemistry and Biochemistry of the Czech Academy of Sciences, Flemingovo nam. 2, 166 10 Prague 6, Czech Republic. ^2^Institute of Physics, Polish Academy of Sciences, Al. Lotnikow 32/46, 02-668 Warsaw, Poland.

