## Supplementary Information for "Coordination of transporter, cargo, and membrane properties during non-vesicular lipid transport"

**Supporting Information**

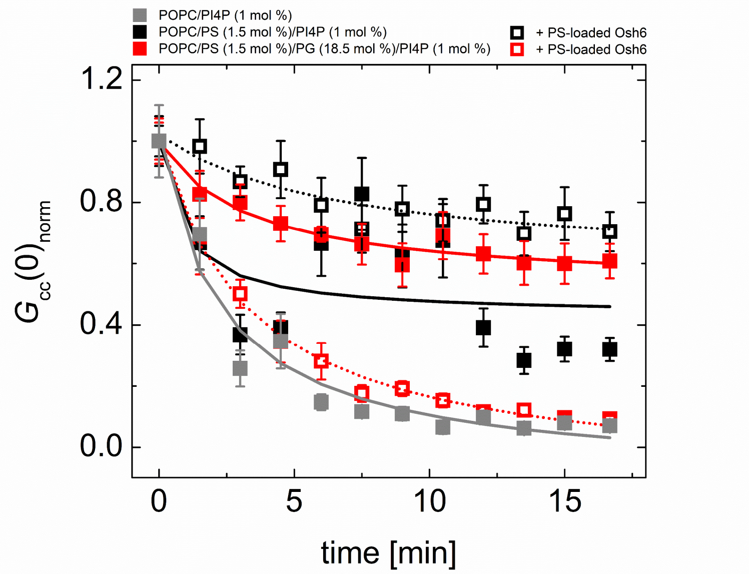


Figure S1. Synchronization of charge and cargo occupancy required for the PI4P extraction from the PM-like membrane. PI4P extraction assay. Temporal dependence of the normalized FCCS read-out, $G_{cc}(0)_{norm}$, monitoring the PI4P drop upon Osh6 addition. PI4P is extracted from various kinds of LUVs: POPC/POPS (1.5 mol %)/PI4P (1 mol %) (black squares), POPC/POPS (1.5 mol %)/ POPG (18.5 mol %)/PI4P (1 mol %) (red squares), and POPC/PI4P (1 mol %) (grey squares). The solid and the hollow squares stand for empty and PS-loaded Osh6 addition, respectively.
